# Use of dual electron probes reveals role of ferritin in erythropoiesis

**DOI:** 10.1101/866889

**Authors:** Maria A. Aronova, Seung-Jae Noh, Guofeng Zhang, Colleen Byrnes, Emily Riehm Meier, Young C. Kim, Richard D. Leapman

## Abstract

Much is known about the finely regulated process of mammalian erythropoiesis that occurs in the bone marrow, whereby erythropoietic stem cells undergo terminal differentiation accompanied by enormous morphological changes to generate highly functional specialized red blood cells. However, a crucial step in erythropoiesis, the labile iron pool and its transport to mitochondria for heme production, is not well understood^1^. We apply a dual 3D imaging and spectroscopic technique, based on scanned electron probes, to measure distributions of ferritin iron-storage protein in *ex vivo* human erythropoietic stem cells, and to determine how those distributions change during terminal differentiation. After seven days of differentiation, the cells display a highly specialized architecture of organelles with anchored clustering of mitochondria and massive accumulation of Fe^3+^ in loaded ferritin cores localized to lysosomal storage depots, providing an iron source for heme production. Macrophages are not present in our *ex vivo* cultures, so they cannot be the source of the ferritin^2^. We suggest that lysosomal iron depots are required by developing reticulocytes while terminally differentiating and continuing to produce heme and globin, which assemble and concentrate to fill the cytoplasm after much of the cellular machinery is expelled.

---

Eukaryotic cellular function depends not only on biochemical reactions of protein assemblies occurring at sub-nanometer scale over times of nanoseconds to milliseconds, but also on ultrastructural alterations of organelles on scales of tens to hundreds of nanometers and over times ranging from minutes to weeks^3,4^. Ultrastructural changes underlie complex processes like endocytosis, intracellular trafficking and differentiation. Sometimes connections between biochemistry and ultrastructure are provided with light microscopy (LM) by identifying specific fluorescently tagged proteins, or with immuno-electron microscopy (EM) by detecting Au-tagged antibodies. However, such labeling schemes are unavailable for tracking specific elements. Here, we show that electron energy loss spectroscopic (EELS)^5-7^ imaging in the scanning transmission EM (STEM)^8,9^ combined with serial block-face scanning EM (SB-SEM)^10^ (Fig. 1, Suppl. Fig. 5) enables quantitative analysis of Fe^3+^ iron within specific organelles of cultured human hematopoietic CD34+ stem cells^11,12^ that are undergoing erythropoiesis^13-15^, the process whereby stem cells terminally differentiate to red blood cells (RBCs) *in vivo*, or to reticulocytes with the *ex vivo* cell cultures in the present study. By sampling cells at different times in their path, from progenitor cells at *t*_*1*_ = 0, through *t*_*2*_ = 7 days and *t*_*3*_ = 14 days, to reticulocytes at *t*_*4*_ = 21 days, we investigate the precisely orchestrated process whereby iron, taken up by the cell, finds its way into mitochondria to enable synthesis of heme, which is eventually incorporated into hemoglobin in the cytoplasm. Erythroblast differentiation *ex vivo* mimics the *in vivo* process, in which progenitor cells accumulate hemoglobin, eventually losing their nuclei and other organelles to form RBCs that fulfill their primary function of oxygen transport through the vascular system.

**Fig. 1.**
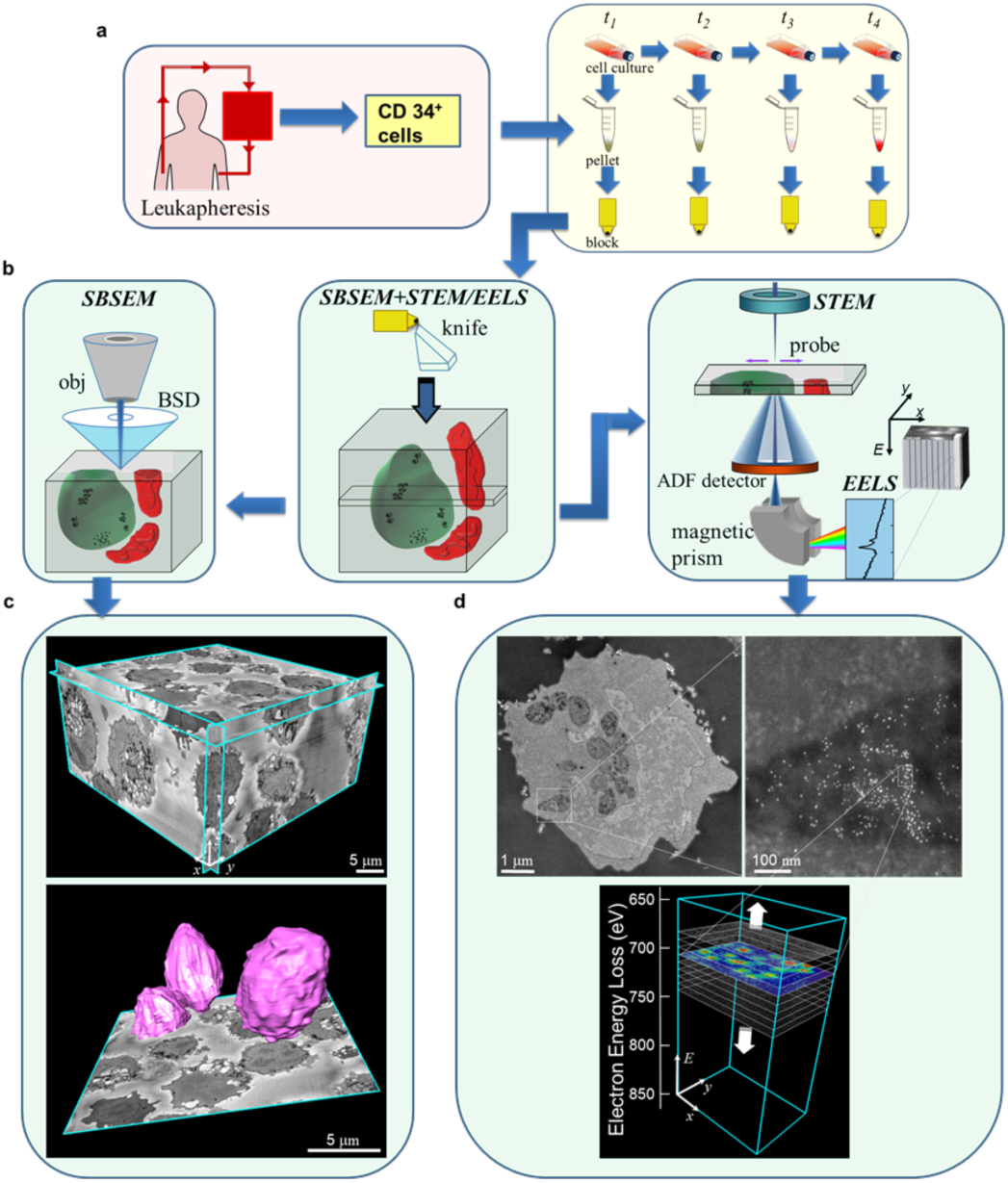
Workflow: sample collection/preparation, scanned probe set-up SB-SEM/STEM, and image analysis. **a**, CD34^+^ stem cells are isolated from healthy volunteers, via leukapheresis, and cultured in serum-free media. At 4 different time points of differentiation: *t*_*1*_ = day 0, *t*_*2*_ = day 7, *t*_*3*_ = day 14 (7 days after erythropoietin (EPO) exposure), *t*_*4*_ = day 19-21 (14 days after EPO exposure), cells are collected and pelleted, to undergo staining and embedding in preparation for electron microscopy (EM); staining protocols described in Methods. **b**, Dual electron beam techniques used to track morphological changes and Fe accumulation during erythropoiesis. (1eft) Serial block-face scanning electron microscopy (SB-SEM): surface of plastic block containing embedded stained cells is scanned with focused electron probe, and image collected with signal from backscattered electron detector; thin section is then removed from block with a diamond knife and block surface re-imaged. This process is repeated until a stack containing hundreds of images is acquired. After image alignment, segmentation provides visualization and quantitation of organelle volumes. (right) Scanning transmission electron microscopy (STEM): thin sections of embedded cells are deposited on TEM grids and imaged by using elastically scattered signal collected with an annular dark-field (ADF) detector, which reveals locations of heavy elements, or by using inelastically scattered signal collected with an electron energy loss spectrometer (EELS) to obtain compositional information at a nanometer scale. **c**, Example of a 3D volume acquired with SB-SEM showing tens of cells at the *t*_*3*_ developmental stage, which enables visualization and quantitation of organelle volumes. **d**, Example of images acquired with ADF-STEM and EELS, showing the cross section of cell with distribution of ferritins, and identification of Fe (represented in color).

It is known that mitochondria play a crucial role in supplying energy and massive levels of heme^16-18^. Fluorescence and histochemical LM microscopy reveal that immature erythroblasts develop an organellar superstructure of mitochondria^19-21^surrounding iron-containing endolysosomes at the Golgi pole (Suppl. Figs. 1, 2, Suppl. Notes 1, 2). To interpret this dynamic morphology shift and its contribution to iron accumulation, requires higher resolution imaging techniques (Fig.1). Although existence of vesicles surrounding mitochondria in erythropoiesis has been reported at the ultrastructural level^22-24^, until now there has been no quantitative analysis of Fe^3+^ transport.

**Fig. 2.**
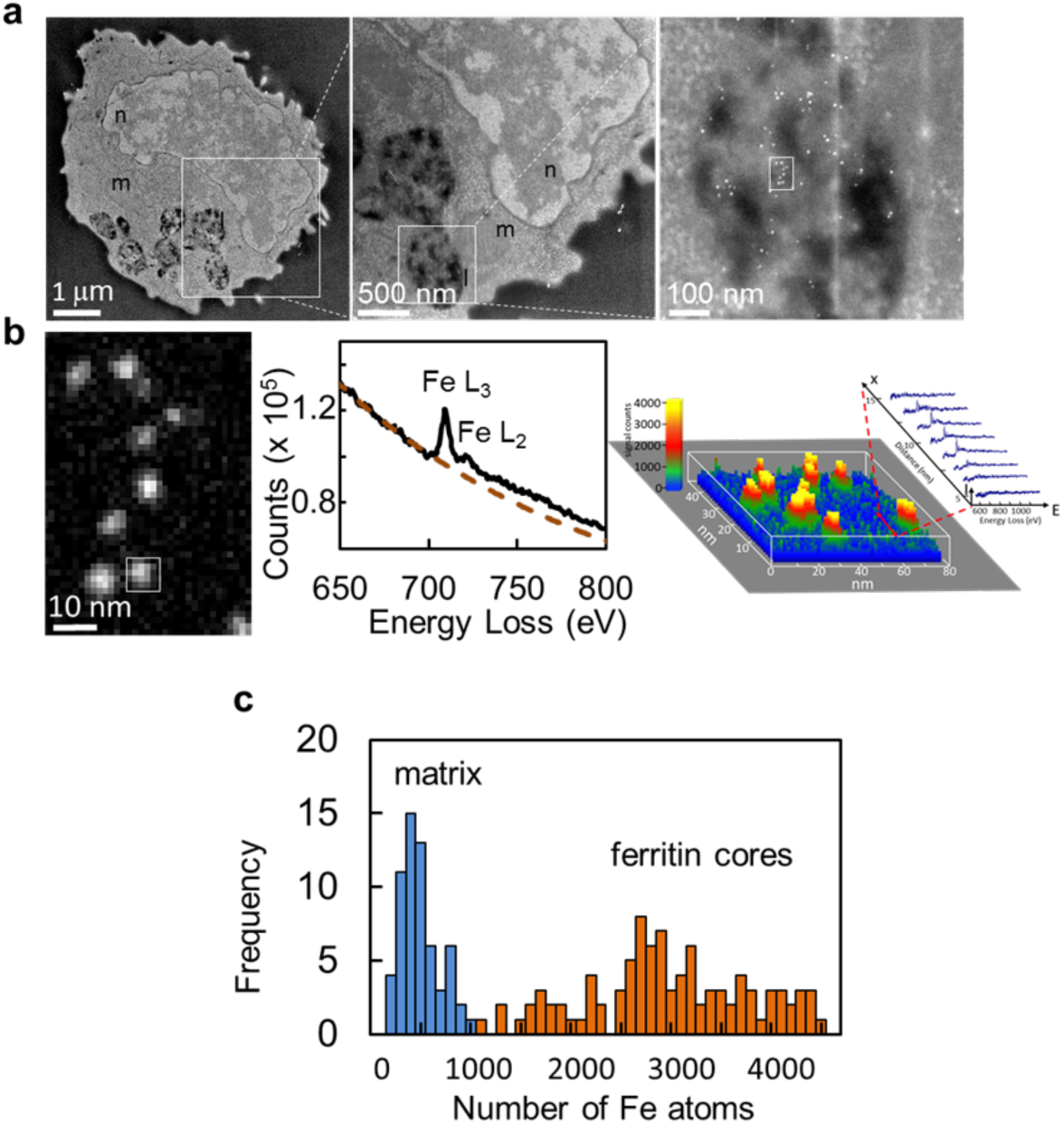
Imaging ferritin distributions in thin sections of differentiating erythroblasts with STEM/EELS. **a**, ADF STEM images illustrated for a *t*_*2*_ erythroblast: whole cell (left) with mitochondria (m) surrounding lysosomes (l), and invaginated by nucleus (n); magnified view (middle) shows proximity of lysosomes and mitochondrion; further magnification (right) shows identifiable individual ferritin cores inside lysosome. **b**, EELS Fe mapping of small rectangular region, outlined in **a** (right panel), shows presence of ferritin cores. In Fe map (left), contrast corresponds to different numbers of Fe atoms in ferritin cores. EELS spectra (middle panel) extracted and averaged over 6 × 6 pixels of one ferritin particle (white box), shows peak at 710 eV corresponding to Fe L_3_ and Fe L_2_ edge, a signature of iron. Surface plot represented in color according to Fe signal (right) in region of ferritin particles and STEM/EELS data as function of *x* (position along sample), *E* (energy loss) and *I* (intensity, counts). As scan line is averaged over 6 pixels (in *y*) and scanned (along *x*, red double headed arrow) from top to bottom of region containing a ferritin particle, the spectral shape in vicinity of Fe edge at 710 eV changes in intensity, depending on the Fe content. **c**, Fe distribution of individual ferritin cores found in lysosomes in 40 cells pooled for *t*_*1*_ – *t*_*4*_ developmental stages together with the surrounding background measurements from other cellular regions. The average Fe content in the ferritin cores was found to be 2,429±817 atoms and average in the surrounding background regions was −59±195 atoms.

STEM and LM images reveal ultrastructural changes in CD34+ cells as they differentiate from *t*_*1*_ to *t*_*4*_ (Suppl. Fig. 2a, Suppl. Fig. 6a), during which, mitochondria and membrane-bound lysosomes gather at the Golgi pole. Within compartments of lysosomes, bright particles ∼ 10 nm in diameter are evident (Fig.2a, Suppl. Fig.6), indicative of ferritin, which is supported by the colocalization of ferritin light chain with the lysosomal protein LAMP1 in fluorescence optical images (Suppl. Fig. 2). We obtain STEM-EELS maps^25^ around the Fe L_2,3_ edge resonance at ∼710 eV, confirming the presence of Fe^3+^ iron in each nanoparticle,^26^ (Fig. 2b, left panel) as illustrated in the spectrum integrated over a single particle (center panel); variation in iron signal across different particles is also evident (right panel). Quantitative EELS analysis of 96 iron-containing particles (Fig. 2c) gives a mean Fe content of 2,430±820 (±s.d.) atoms, consistent with the known iron content of isolated ferritin cores^27,28^ (Suppl. Fig. 3, Suppl. Note 3). Then, by counting numbers of particles per unit area in STEM images from lysosomes and knowing the section thickness (∼100 nm), we can estimate the number of ferritin cores, and therefore the number of iron atoms per unit volume of lysosomes. Then, from SB-SEM data, we can measure the volume of lysosomes per cell and accordingly the total number ferritin iron atoms per cell (Suppl. Note 4, Suppl. Fig. 7).

**Fig. 3:**
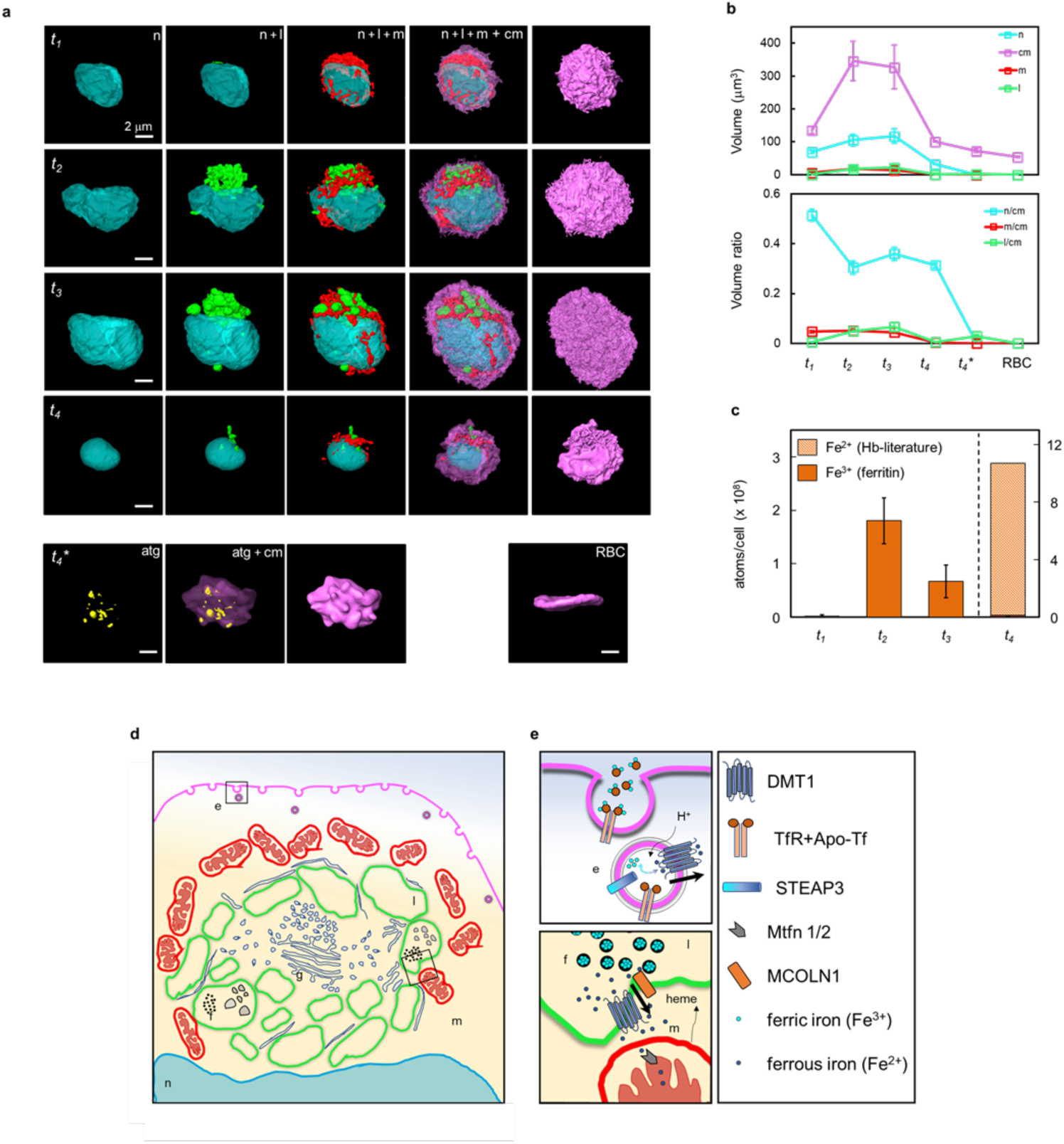
Morphological analysis of differentiating erythroblasts by SB-SEM reveals changes in organelle volumes and evolution of Fe3+ content; a model of cellular processes involved in erythropoiesis. **a**, Surface rendering of SB-SEM segmented data from representative cell at each development stage, *t*_*1*_, *t*_*2*_, *t*_*3*_ and *t*_*4*_ depicting evolution of cell morphology during erythroblast differentiation; cyan corresponds to nucleus (n), green-lysosomes (l), red-mitochondria (m), magenta-cell membrane (cm), and yellow-autophagosomes (atg). Lower row of images show: *t*_4_^*^ – reticulocyte without nucleus, which resembles RBC (shown for comparison at right) but with bumpy plasma membrane. **b**, Volume and volume fractions calculated from segmentation of each cellular development stage; standard errors of the mean from 5 different cells analyzed at each developmental stage are indicated (data shown in Supplemental Fig. 4). **c**, Measured ferritin Fe^3+^ in atoms per cell as function of development stage (solid bars) together with Fe^2+^ content of hemoglobin (lighter bar) in atoms per *t*_*4*_-reticulocyte, based on same concentration and volume of hemoglobin as in RBC. **d, e** Proposed model describing iron trafficking during erythropoiesis based on existing knowledge and current findings. **e**, Transferrin receptor (TfR)-mediated endocytosis is utilized to capture extracellular ferric iron (Fe^3+^) by binding to apo-transferrin (apo-Tf); intra-endosomal pH is lowered in endosomal vesicles, enabling dissociation of iron from transferrin and reduction of Fe^3+^ to Fe^2+^ through metalloreductase enzyme STEAP3; reduced Fe^2+^ iron exits endolysosomes via divalent metal transporter (DMT1) into cytoplasm where it combines with Golgi-produced apo-ferritin shells and is oxidized back to Fe^3+^ to form crystalline ferritin cores. Subsequently, ferritin molecules together with other small vesicular bodies are engulfed by membranes via autophagosomal pathway into large lysosomes (l). Our hypothesis suggests that Fe^2+^ is trafficked via DMT1 or mucolipin-1 (MCOLN1) to mitochondria for inclusion into heme and iron-sulfur clusters through incompletely understood mechanisms.

3D images of erythroblasts were segmented to obtain volumes of entire cells, nuclei, mitochondria, and lysosomes for 5 cells at each stage of development (Fig. 3a, Suppl. Fig. 4, Supp. Table 1, Suppl. Movie 2). Compartment volumes increase from *t*_*1*_ to *t*_*2*_, remain relatively constant up to *t*_*3*_, and then decrease at *t*_*4*_ (Fig. 3b). The nuclear volume almost doubles from *t*_*1*_ to *t*_*2*_, elongating and developing invaginations to accommodate lysosome formation at the Golgi pole where mitochondria gather (Fig. 3a, Suppl. Fig 2a, Suppl. Fig. 4). At *t*_*4*_, nuclei become spherical with volumes fourfold lower than at *t*_*3*_, before ejection. At *t*_*1*_, few endolysosomes are evident, whereas at *t*_*2*_ and *t*_*3*_ the lysosomal volume increases more than 30-fold to accommodate iron storage. Finally, at *t*_*4*_, organelles are digested via the autophagosomal pathway. Mitochondrial networks in close association with lysosomes mainly aggregate at the Golgi pole with formation of a side ridge. The mitochondrial volume increases twofold from *t*_*1*_ to *t*_*3*_, to accommodate heme production, after which it decreases, eventually being ejected through autophagosomal digestion. The cell membrane is highly ruffled at *t*_*1*_, *t*_*2*_ and *t*_*3*_, and becomes smooth only after organelles are digested and nucleus extruded. We infer, that this membrane morphology, as well as a threefold increase in cell volume from *t*_*1*_ to *t*_*3*_, provides increased surface area for docking of additional transferrin receptors carrying iron for heme production. Despite enucleation at *t*_*4*_, terminally differentiated erythroblasts in *ex vivo* cultures resemble reticulocytes and do not form smooth biconcave discs characteristic of RBCs, since they must enter the blood stream *in vivo* to fully mature (Fig. 3a, lower panel). Nevertheless, our estimates of reticulocyte heme concentration match the known value for RBCs (Suppl. Fig. 1c). Throughout *t*_*1*_, *t*_*2*_, and *t*_*3*_ differentiation, cellular machinery must continue to produce massive amounts of heme and globin, even while the cell reorganizes.

Combining STEM-EELS with SB-SEM data, we now estimate numbers of Fe^3+^ iron atoms within ferritin cores per cell at different stages of development (Fig 3c). Furthermore, we relate numbers of Fe^3+^ iron atoms to heme production, which starts prior to *t*_*3*_ and increases threefold at *t*_*4*_ (Suppl. Fig. 1c). Our results show that most ferritin accumulation occurs in lysosomes and, toward later stages of development, ferritins aggregate into clusters (Suppl. Fig. 6). From quantitative electron microscopy (Suppl. Note 4, Suppl. Fig. 7, Suppl. Fig. 9, Fig. 3c) and the analytical biochemical analysis (Suppl. Fig 1c), we deduce, that the iron stored in ferritin reaches a maximum at *t*_*2*_ and then decreases through continuing stages of differentiation as heme and hemoglobin accumulate, reaching a plateau at t*4*, which closely resembles the state of mature RBCs. Continuous production of heme and hemoglobin during cell divisions after addition of erythropoietin in erythropoiesis is well established^12^, and it is known that loss of surface transferrin receptors does not occur until a state of terminal maturation is reached^29^. Analysis of STEM tomograms and observation of a tight 5-10 nm apposition between mitochondrial and lysosomal membranes (Suppl. Fig. 8, Suppl. Mov. 1), strongly supports the hypothesis that the transfer of Fe^2+^ occurs from ferritin stored in the lysosomes^30^ to mitochondria.

In our model for iron uptake during erythroblast differentiation (Fig. 3d and 3e), based on the present and previous work^31-39^, transferrin-mediated endocytosis first captures extracellular Fe^3+^ into endosomal vesicles, whose pH is then lowered, allowing dissociation of Fe^3+^ from transferrin, and reduction to Fe^2+^. Reduced Fe^2+^ exits endolysosomes to the cytoplasm where it combines with apoferritin. Fe^2+^ is then oxidized to Fe^3+^ in the ferritin cores, before being enclosed in larger lysosomal vesicles. Fe^3+^ is eventually converted back to Fe^2+^ and trafficked to mitochondria, which surround the lysosomes to enable heme production. This model suggests that dynamic morphological changes involving mitochondria, lysosomes, Golgi, nuclear membrane, and plasma membrane, are crucial for maturation of cultured reticulocytes. As erythroblasts undergo terminal differentiation to reticulocytes, composed essentially of highly concentrated hemoglobin, they must continue to produce globin and heme in a milieu where much of the cellular machinery becomes increasingly impeded. We suggest, that only critical portions of the machinery remain, including ribosomes and mRNA for globin synthesis, mitochondria for heme synthesis, ATP molecules for energy, and iron storage by ferritin holoproteins sequestered in lysosomes, whose membranes eventually also function to expel the residual organelles from the cell. Our observations support an intimate relationship between erythropoiesis and iron homeostasis in differentiating erythroblasts, and we suggest a similar process occurs *in vivo* in maturing RBCs within bone marrow, although on a faster timescale^40^.

Our results suggest a function for biology’s natural hybrid nanoparticle, ferritin, in erythropoiesis, which in humans produces 2,000,000 new red blood cells each second. Perhaps this hitherto unknown role is not surprising since approximately 25% of the body’s iron is stored as ferritin, which is ubiquitous in every cell. A more complete understanding of iron processing at the subcellular level could elucidate *in vivo* erythropoiesis taking place in erythropoietic islands of bone marrow^41^, as well as improve our ability to manufacture red blood cells in *ex vivo* cultures, which could advance transfusion medicine^42^.

## Acknowledgments

The Intramural Research Program of the National Institute of Biomedical Imaging and Bioengineering, the National Institute of Diabetes and Digestive and Kidney Diseases, and Office of Naval Research via the U.S. Naval Research Laboratory base program supported this work. We thank Dr. J. L. Miller for discussions on the topic of erythropoiesis.

## Author Contributions

RDL and MAA conceived and directed the project. MAA and SJN performed the experiments and analyzed the data. GZ prepared all electron microscopy specimens for both SB-SEM and STEM/EELS analysis. CB established primary cell culture and conducted flow cytometry analyses. ERM performed HPLC. YCK wrote and implemented MATLAB code to calculate atomic distributions. RDL, MAA and SJN wrote and revised the final manuscript.

## Additional information

Supplementary information is available for this paper with initial submission. Correspondence and requests for materials should be addressed to MAA or RDL

## Methods

### Cell cultures

All studies involving human subjects were approved by the institutional review boards of the National Institute of Diabetes, Digestive, and Kidney Diseases. After informed consent was obtained, human CD34(+) cells were collected from peripheral blood of healthy volunteers at the National Institutes of Health (Bethesda, MD). Using primary human CD34+ stem cells, which were isolated via Leukapheresis (Fig 1a) and cultured *ex vivo* for 21 days, four cell populations were isolated: *t*_*1*_[CD34(+)], *t*_*2*_ [CD36(+), CD235(-) erythroblast], *t*_*3*_ [CD36(+), CD235(+) erythroblast], and *t*_*4*_ (nucleated and enucleated erythrocytes)^11,12^. Overall, primary erythroblasts in culture demonstrated well-coordinated cellular events including transferrin receptor upregulation, and increased heme and hemoglobin production that are largely mediated by erythropoietin (Suppl. Note 1, Suppl. Fig. 1). Cell culture and differentiation assessments were previously described^11,12^.

### Reagents and antibodies

The following reagents/primary antibodies were used for immunofluorescence analyses: paraformaldehyde solution (10%) was purchased from Electron Microscopy Sciences (Hatfield, PA); mouse anti-TfR2, rabbit anti-DMT1 (Santa Cruz Biotechnology, Santa Cruz, CA)); rabbit anti-lysosomal-associated membrane protein 1 (LAMP1), rabbit anti-Giantin, rabbit anti-γ-tubulin (Abcam, Cambridge, MA); mouse anti-Rab11 (Millipore, Billerica, MA); mouse anti-DDK (4C5) (OriGene Technologies, Rockville, MD). Alexa-594 labeled transferrin, Mitotracker Red, and Alexa-488 or Alexa-594 labeled anti-mouse IgG or anti-rabbit IgG secondary antibodies were from Invitrogen (Carlsbad, CA). Mounting medium for fluorescence with DAPI staining was purchased from Vector Laboratories (Burlingame, CA).

### Sulfide-silver method (iron staining)

Initial studies of intracellular iron were performed by sulfide-silver staining as previously described^43^. All chemicals employed in sulfide-silver staining were obtained from Sigma Aldrich. In brief, cells were fixed with 2% glutaraldehyde in 0.1 M Nacacodylate (pH 7.2) with 0.1 M sucrose for up to 2 hours at room temperature. After washing 5 times with H_2_O, samples were sulphidated with 1% ammonium sulphide (pH ∼9) in 70% ethanol for 15 min then washed with H_2_O. Heavy metals (mainly iron in cells) were visualized by autometallography in a colloid-protected developer containing silver nitrate and hydroquinone in citrate buffer in the dark for 20 to 30 minutes.

### Measurement of heme and hemoglobin

The QuantiChrom heme Assay (BioAssay Systems) was used to measure heme in cultured erythroid cell lysates (*t*_*1*_ *- t*_*4*_). The amount of heme in each sample was normalized per microgram of total protein. Hemoglobin in erythroid cells at *t*_*2*_ and *t*_*4*_ (2 million cells each) was detected by HPLC as previously described^44^.

### Electron microscopy, SB-SEM

From cultured cells we took 4 developmental stages: *t*_*1*_ = day 0, *t*_*2*_ = day 7, *t*_*3*_ = day 14 (7 days after erythropoietin (EPO) exposure), *t*_*4*_ = day 19-21 (14 days after EPO exposure). Combinatorial heavy metal staining protocol was followed, developed by Mark Ellisman’s group^45,46^. The cells were fixed for five minutes in a mixture of 2.5% glutaraldehyde and 2% formaldehyde in sodium cacodylate buffer with 2 mM calcium chloride. Then they were fixed for another two to three hours on ice in the same solution, after which they were thrice rinsed (2x or 3x) with cold cacodylate buffer containing 2 mM calcium chloride (five minutes for each rinse). Next came heavy metal staining: the cells were fixed in Reduced Osmium—a solution containing 3% potassium ferrocyanide in 0.3 M cacodylate buffer with 4 mM calcium chloride combined with an equal volume of 4% aqueous osmium tetroxide—for one hour on ice. They were then placed in a 0.22 μm- Millipore-filtered 1% thiocarbohydrazide (TCH) solution in ddH_2_O for 20 minutes. The heavy metal staining phase concluded with fixation of the samples in 2% osmium tetroxide in ddH_2_O for 30 minutes. Between each of the preceding steps, the samples were washed five times, three minutes each time, with ddH_2_O. At last, the samples were placed in 1% aqueous uranyl acetate and left overnight at ∼4°. The next day, after once again washing the samples with ddH_2_O five times for three minutes each time, we performed en bloc Walton’s lead aspartate staining^47^: the cells were submerged in a lead aspartate solution and placed in the oven for 30 minutes. Then, following another five three-minute ddH_2_O rinses, the samples were dehydrated and embedded in Epon-Araldite according to standard protocols. Our final sample preparation step was to mount the islets for SB-SEM with the goal of minimizing specimen charging. Each resin-embedded sample was first mounted on an empty resin block to be trimmed under the microtome. Once its top was exposed, each block was re-mounted, exposed side down, to a special aluminum specimen pin (Gatan, Pleasanton, CA) using Circuit Works Conductive Epoxy (CW2400); CW2400 served to electrically ground the cells to the aluminum pin. The re-mounted samples were then trimmed again, and, finally, sputter-coated with a thin ∼4 nm layer of gold.

The trimmed resin-embedded stained cells were imaged using a Gatan 3View serial block-face imaging system installed on a Zeiss SIGMA-VP SEM, operating at an accelerating voltage of 2.7 kV using a 60 μm condenser aperture. The SEM was operated at 25 Pa gas pressure. The acquired images had a pixel size of 7.7 nm in the x-y plane and 50 nm along the z-axis. The resulting datasets were assembled into volume files and aligned using Digital Micrograph (Gatan, Inc.), binned by 4, and then manually segmented into 3-D models in Amira (FEI, Thermo Fisher Scientific). Some thresholding was used to expedite the process. The volumes of the segmented organelles: mitochondria, nuclei, vesicles and cell membrane were measured by Amira software’s “Material Statistics” module, thus obtaining all the reported three-dimensional size parameters. Measurements were obtained from 27 cell membranes, 21 nuclei, 20 mitochondrial networks, and 25 vesicle networks, with 5 cells selected from each of the 4 developmental stages. All cell-type differentiation was performed by visual inspection, and cell types were assessed according to established morphologies^48^.

### Electron microscopy, STEM/ EELS

Cells from different time stages were pelleted and fixed in a mixture of 2.5% formaldehyde and 2.0% glutaraldehyde in PBS (pH= 7.4), followed by 0.02% osmium tetroxide fixation. After several rinses in the buffer (3×10min), the samples were dehydrated in a series of ethanol (20%, 40%, 60%, 75%, 95% for 10 min and 100% for 30 min with 3 changes) and infiltrated with Epon-Aradite (Ted Pella, Redding, CA) for 2 days (25% of Epon-Aradite and ethanol for 2h, 50% for 4h, 75% for 8h and 100% for 1 day with 2 changes). The samples were polymerized at 60°C for 2 days.

Other samples were prepared by fixing cells in a mixture of 2.5% formaldehyde and 2.0% glutaraldehyde in PBS, briefly rinsed in 0.1 M sodium cacodylate buffer, and postfixed in 1.0% osmium tetroxide plus 0.8% potassium ferricyanide in the same buffer for 60 min. After several rinses in PBS (3x for 10 min), the samples were dehydrated in an ethanol gradient and infiltrated with Epon-Aradite (Ted Pella, Redding, CA) for a couple days. The samples were then polymerized at 60°C for another 2 days. For all different time stages the samples were prepared and imaged the same way. 100-120 nm sections were deposited on TEM copper girds.

To compare results with a pure ferritin-standard sample, horse spleen ferritin (Sigma-Aldrich, US) was deposited on a thin carbon film (∼5 nm) and was let to dry Dark field STEM and STEM/EELS images were acquired using Tecnai TF30 electron microscope (FEI, Thermo Fisher Scientific), equipped with a Quantum imaging filter (Gatan Inc., Warrendale, PA), and operating at an accelerating voltage of 300 kV. The energy loss range was 445 eV-1470 eV, and the elemental maps were taken from a small area of the specimen, using drift correction when needed. Elemental images containing about 50 by 50 pixels were acquired with a pixel size of 1-2 nm.

To determine the iron loading in the ferritin particles (standard sample) after imaging with STEM/EELS, the spectral tool in the Digital Micrograph software (Gatan Inc.) was used to select individual ferritin particles. The background under the Fe L_2,3_ core edge at 710 eV was removed by fitting the inverse power law^5^ and the Fe L_2,3_ signal was summed over a 10-eV window, to generate an integrated Fe signal map. The iron map was quantified calculated by dividing the net Fe signal by the inelastic cross section and by the total beam current. Thus, the number of Fe atoms *N*_*Fe*_ in each analyzed ferritin particle was be calculated according to the following 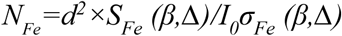, where *S*_*Fe*_ is signal (number of counts in the Fe L edge), *I*_*0*_ is the total incoming current (number of counts in the zero loss and low-loss spectrum), *σ*_*Fe*_ is the inelastic cross section, and *d* is the pixel size in nm. *β* is the collection semi-angle defined by the spectrometer entrance aperture, which was 20 mrad and *Δ* is the integration energy window, which was set to be 10 eV in order to include the L_3_ white line resonance. A value for the scattering cross section *σ*_*Fe*_ is not well established with a wide range of values reported in previous literature. We have therefore calculated the L_3_ white line cross section both from x-ray data and from empirical EELS measurements on nanoparticles of known composition (Suppl. Note 3). The value used here for *σ*_*Fe*_ *(β, Δ)* was estimated to be 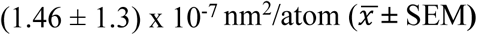.

## Data availability

The data that support the findings of this study are available from the corresponding authors upon request.

