## Supplementary Information for "Use of dual electron probes reveals role of ferritin in erythropoiesis"

*Corresponding Authors

**Supplementary Note 1**

In vertebrate cells transferrin-mediated endocytosis relays iron to the mitochondria for vital cellular requirements and heme production[49]. Even though the presence of the transferrin receptor (CD71) expression was detected prior to addition of erythropoietin (EPO), there was a sharp increase in CD71 expression on the plasma membrane with the addition of EPO after $t_2$ (Suppl. Fig. 1a). The loss of CD71 expression on the cell surface during the final stages of differentiation is morphologically associated with nuclear condensation and loss of the perinuclear vesicular body. The vesicular body is prominent in less mature cells that are morphologically similar to circulating leukemic erythroblasts[50]. With differentiation, the vesicular body shrinks and is eventually lost. Negligible heme and hemoglobin were detected prior to EPO (Suppl. Fig. 1a, c). After the addition of EPO, the heme and hemoglobin levels rose rapidly. In the absence of EPO, transferrin receptor expression is not fully upregulated, neither heme nor hemoglobin accumulates, and the immature erythroblasts undergo apoptosis (data not shown).

For initial localization of iron in the cells, erythroblasts were counterstained with silver sulfide resulting in detection of iron through nucleation of silver deposits visible as dark brown pigments (Suppl. Fig 1b). While not detected in the undifferentiated $t_1$ cells, iron-containing precipitates were present in the vesicular superstructures during the later stages of differentiation as the cells accumulate hemoglobin prior to enucleation. For comparison, iron precipitates were neither detected in erythroblasts cultured in apo-transferrin, nor in non-erythroblast cells.

### Supplementary Figure 1

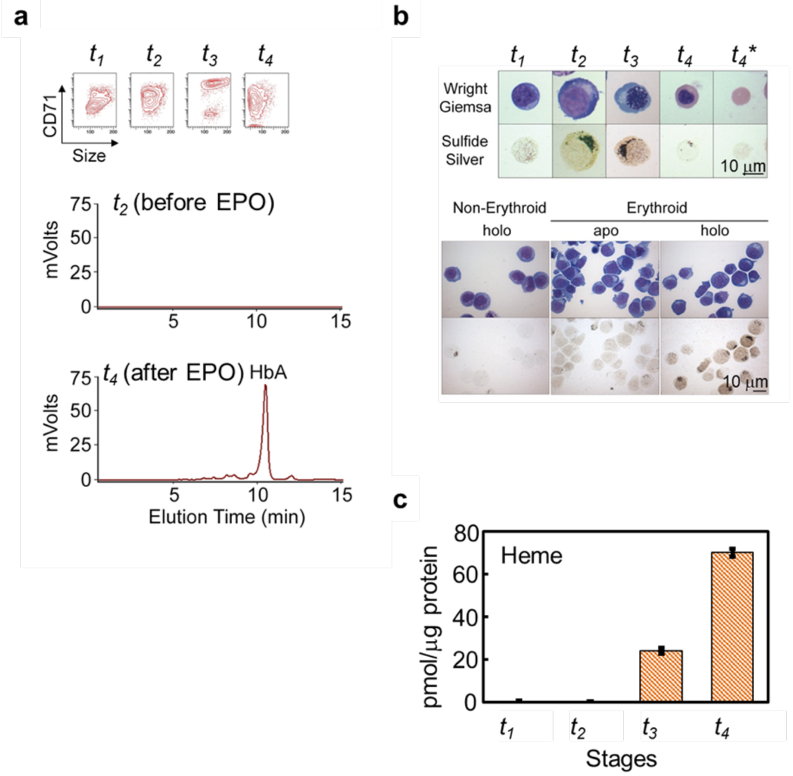


**Suppl. Figure 1.** Erythroblast iron localization. (a) Flow cytometric profiles, in which the y-axes on the flow contour plots represents transferrin receptor (CD71) and the x-axis, the size (forward light scatter). Initially, there is gain and then gradual loss of transferrin receptor expression during erythropoiesis for $t_1$-$t_4$ time periods. HPLC measurements of hemoglobin (HbA) at $t_2$ (pre-EPO) and at the final stage of differentiation at t4. (b) Wright-Giemsa staining of erythroblasts counterstained with sulfide-silver method (SSM) for iron deposit detection. The iron is viewed as silver precipitates, which are the dark brown pigments. At $t_2$ stage the cells cultured in holo-transferrin-$Fe^{3+}$ bound (0.1 mg/ml) and apo-transferrin (0.1 mg/ml) were sorted by FACS into erythroid/and non-erythroid lineage and stained with SSM. The silver deposits are detected in the erythroid lineage in media supplemented with holo-transferrin. Note: $t_4$* corresponds to t4 stage without the nucleus. (c) Cellular heme measurements during $t_1$-$t_4$ culture period. Heme was measured as a portion of the total protein content (pmol/μg protein). All error bars indicate standard deviation (SD).

### Supplementary Note 2

It is evident that mitochondria become organized around one pole of the immature cells and surround the clustered lysosomes. In addition, ferritin (FTL-ferritin light chain) is colocalized with the lysosomes (LAMP1) again surrounded by mitochondria (Suppl. Fig. 2b). Transferrin receptor 2 (TfR2) was also colocalized with the lysosomes and partially colocalized with divalent metal transporter (DMT1) and recycling endosomal marker (RAB11), as was the centrosomal marker γ-tubulin (Suppl. Fig 2c). TfR2 did not co-localize with giantin, a Golgi marker, although Golgi was found in this same region, but more proximal to the nucleus.

### Supplementary Figure 2

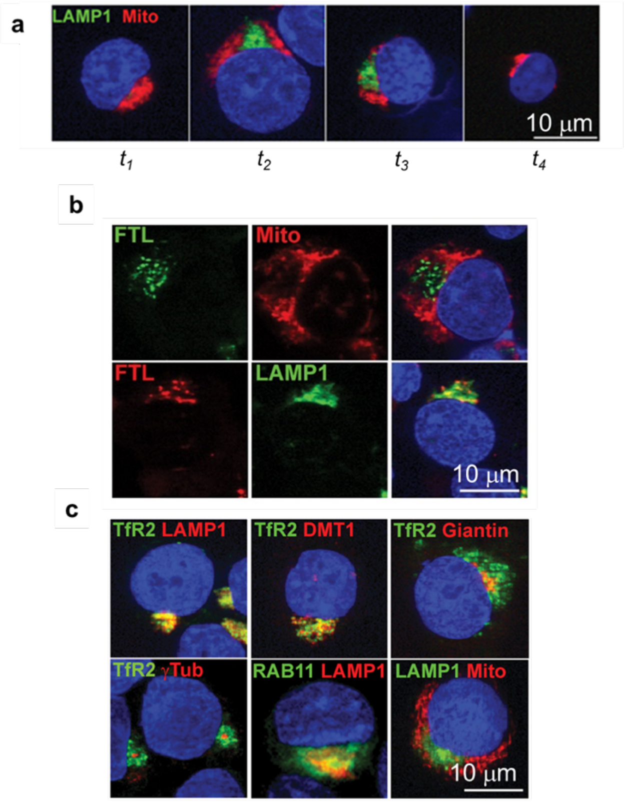


**Suppl. Figure 2.** Confocal imaging of erythroblasts: (a) Cells were stained for mitochondria (Mito, red) and lysosomes (LAMP1, green) at indicated culture stages during erythroid differentiation. Mitochondria and lysosomes are closely related at the pole of the cells. (b) At $t_2$ stage the following was labeled: ferritin light chain (FTL, left panel), mitochondria (Mito, upper center panel), and lysosomes (LAMP1, lower center panel), the merging of FTL with Mito and FTL with LAMP1 are presented in the right panels. This confirmed our hypothesis that ferritins are colocalizing with lysosomes, surrounded by mitochondria at the pole of the cell. (c) Confocal analyses of perinuclear compartments. At $t_3$, the cells were immuno-stained for confocal analyses of the perinuclear compartment with the following antibodies: TfR2, transferrin receptor 2; LAMP1, lysosome-associated membrane protein 1; DMT1- divalent metal transporter1; Giantin-Golgi marker; γ-tubulin, centrosomal marker; RAB11- recycling endosomal marker; Mito-mitochondria.

**Supplementary Note 3**

**A. Calculation of cross section $\sigma_{Fe}$ (β, Δ) for Fe $L_{2,3}$ white line**

The cross section was calculated in two ways.

1. The relation between photoabsorption cross section and the non-relativistic electron energy loss differential cross section is given by[51]

$$\frac{\sigma_\gamma(E)}{\left[\frac{d\sigma(E)}{dE}\right]} = \frac{2\pi a_0}{\hbar c}\frac{E_0 E}{ln\left[\frac{4E_0}{E}\right]}, \qquad \text{(S1)}$$

where $E$ is the photon energy or energy loss, $E_0$ is the primary electron energy, $a_0$ is the Bohr radius, $\hbar$ Plank's constant divided by 2π, and $c$ is the speed of light. And $\frac{d\sigma(E)}{dE}$ is the electron energy loss differential cross section. Using the relativistically corrected beam energy $T_0$ instead of $E_0$, we can write

$$\left[\frac{d\sigma(E)}{dE}\right] = ln\left[\frac{4T_0}{E}\right]\left[\frac{1}{1.68\times 10^{-3}T_0}\right]\frac{\sigma_\gamma(E)}{[E]}, \qquad \text{(S2)}$$

Experimental x-ray absorption data measures absorption coefficient $\mu(E)$ instead of absorption cross section $\sigma_\gamma(E)$:

$$\mu(E) = n\sigma_\gamma(E), \qquad \text{(S3)}$$

where $n$ is the number of atoms of element per unit volume. Now considering the Fe $L_3$ white line excitation at 708±5 eV and integrating the photoabsorption spectrum and the electron energy loss spectrum, for an incident electron energy of $E_0$ = 300 keV or $T_0$ = 154.1 keV[52].
Then,

$$\int \frac{d\sigma(E)}{dE} dE = \frac{0.0262}{nE_{Fe-white\ line}} \int \mu(E) dE, \qquad \text{(S4)}$$

$$n = \frac{\rho_{Fe} N_0}{A_{Fe}}, \qquad \text{(S5)}$$

where $\rho_{Fe}$ is the density of Fe, $N_0$ is the Avogadro's number and $A_{Fe}$ is the atomic weight of Fe. From these constants n is calculated to be 85 Fe atoms/nm$^3$

$$\int \frac{d\sigma(E)}{dE} dE = \frac{0.0262}{n} \int \mu(E) dE = 1.17 \times 10^{-7} nm^2/atom,$$

The integral of $\mu(E)$ is obtained from published x-ray absorption data of Regan T. J., et al.[53] with an estimated uncertainty of 5%. Therefore $\boldsymbol{\sigma_{Fe}}$ **= (1.17 ± 0.06) ×10$^{-7}$ nm$^2$/atom**

2. We calculate $\sigma_{Fe}$ using experimental data from material of a known composition containing Fe: $Fe_2O_3$ and $Fe_3O_4$ nanoparticles. We can write:

$$N_{Fe} = \left(\frac{S_{Fe}(\beta,\Delta)}{I_0 \sigma_{Fe}(\beta,\Delta)}\right) * d^2 \quad N_O = \left(\frac{S_O(\beta,\Delta)}{I_0 \sigma_O(\beta,\Delta)}\right) * d^2, \qquad \text{(S6)}$$

$$\frac{N_O}{N_{Fe}} = \frac{S_O(\beta,\Delta)\, \sigma_{Fe}(\beta,\Delta)}{\sigma_O(\beta,\Delta) S_{Fe}(\beta,\Delta)}, \qquad \text{(S7)}$$

Here, $S_{Fe}$ and $S_O$ are integrated signals for iron and oxygen respectively, $I_0$ is the total incoming electron dose, $\sigma_{Fe}/\sigma_O$ is the inelastic cross section for Fe/O, and $d$ is the pixel size in nm. $\beta$ is the collection semi-angle defined by the spectrometer entrance aperture, which was 20 mrad and $\Delta$ is the integration energy window, which is 10 eV for Fe and 100 eV for oxygen. Knowing the $\sigma_O$ and the ratio of elements found in the material we can solve for $\sigma_{Fe}$.

To find out $\sigma_O$ at beam energy of 300 keV, we use data from Egerton R. F. (1979)[54], for $\sigma_O$ at beam energy of 100 keV = 3.5x10$^{-7}$ nm$^2$/atom and Eq. 2.

$$\frac{\sigma_{O,E=300keV}}{\sigma_{O,E=100keV}} = f,\ f = \frac{T_0(100\ keV)}{T_0(300\ keV)} \frac{ln\left[\frac{4E_0(300keV)}{580}\right]}{ln\left[\frac{4E_0(100keV)}{580}\right]}, \qquad \text{(S8)}$$

Where the calculated $f$ is 0.581, which gives rise to $\sigma_O$ at 300 keV = 2.03 x 10$^{-7}$ nm$^2$/atom

From 23 particles the $\sigma_{Fe}$ = (1.46 ± 0.13) ×10$^{-7}$ nm$^2$/atom, ($\overline{x}$ ± SEM)

### B. Ferritin Test sample

We first conducted a test using horse spleen ferritin particles deposited on the carbon film and imaged them with STEM/EELS. Suppl. Fig 3 shows the workflow of how this was accomplished: from the EELS maps, the signal was calculated and using Eq. 1, including the value of the inelastic cross section that we have calculated as well. The histogram that was plotted, which revealed that on average there are 2,600±842 Fe atoms in a single ferritin core, and 80±70 Fe atoms in the regions of carbon film, devoid of Fe. Applying the same process and analyzing more than 10 cells from each $t_1$-$t_4$ stages (Suppl. Fig 6), a distribution of Fe content in ferritin particles is shown in Figure 3d and has a 2,429±817 Fe atoms, consistent with the known composition of ferritin cores (the area in the matrix was measuring -59±195 Fe atoms).

**Supplementary Figure 3**

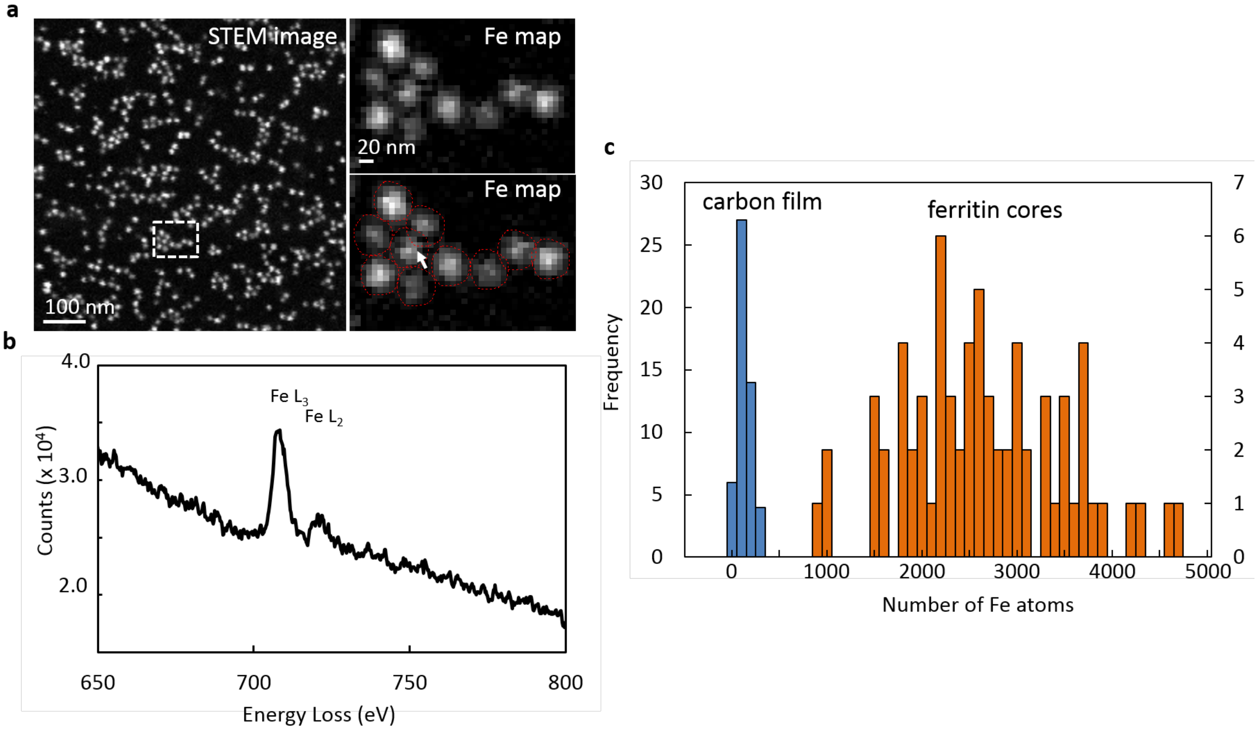


**Suppl. Figure 3.** Quantitative analysis of the Fe distribution in isolated horse spleen ferritin deposited on a thin carbon film. (a) Dark field STEM image of a region with many ferritin particles. EELS data was acquired from a region, outlined with a white dotted line. The Fe map (right top) was generated by integrating the spectral signal (b) at the Fe $L_{2,3}$ edge at 710 eV after background subtraction, incorporating inelastic scattering cross section (see Suppl. Info) and the total electron dose. Then the intensities, in numbers of Fe atoms, were integrated for each ferritin particle (right bottom) to generate a histogram in (c), which shows that a single ferritin core has on average 2,600±842 Fe atoms. (In the carbon film regions, devoid of Fe, the measurements showed 80±70 Fe atoms.)

**Supplementary Figure 4**

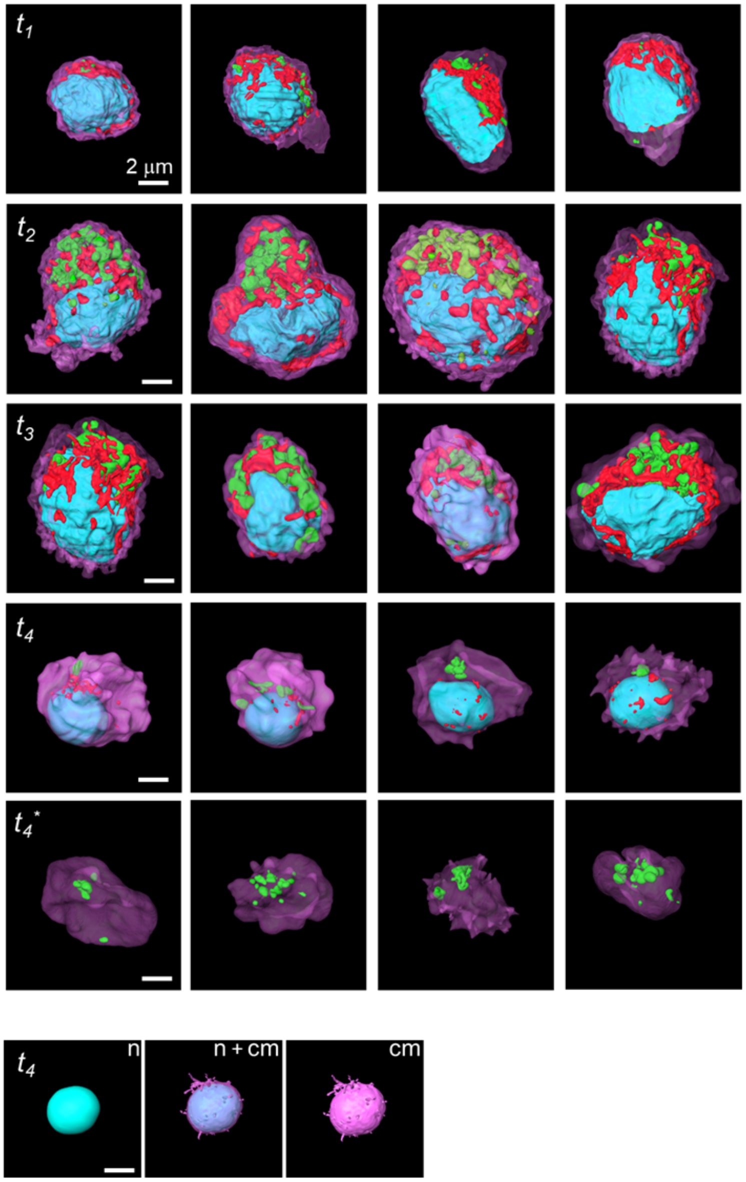


**Suppl. Figure 4.** Complementary segmentation of additional cells for stages $t_1$-$t_4$, which are shown in the main text, Figure 3a, whose calculated volumes and volume ratios together with their respective errors (s.e.m.) were used to generate the graphs in Figure 3b.

**Supplementary Table 1**

Mean volume of nuclei, mitochondria, lysosomes, and cell for different stages of development; and organelle-to-cell volume ratios. These values are plotted in Figure 3b.

Volume ($\mu m^3$)

| | nucleus | s.e.m. | cell | s.e.m. | mitochondria | s.e.m. | lysosome | s.e.m. |
|---|---|---|---|---|---|---|---|---|
| $t_1$ | 69.59 | 5.32 | 135.18 | 11.37 | 6.34 | 0.60 | 0.60 | 0.34 |
| $t_2$ | 105.26 | 17.33 | 345.68 | 60.20 | 17.80 | 4.56 | 17.20 | 3.46 |
| $t_3$ | 117.43 | 21.79 | 327.26 | 66.84 | 14.27 | 3.55 | 21.74 | 4.07 |
| $t_4$ | 31.54 | 1.05 | 100.82 | 2.31 | 0.31 | 0.11 | 0.44 | 0.12 |
| $t_4$* | - | - | 72.19 | 5.90 | - | - | 2.09 | 0.59 |
| RBC | - | - | 54.20 | 2.88 | - | - | - | - |

Volume ratio

| | nucleus/cell | s.e.m. | mitochondria/cell | s.e.m. | lysosome/cell | s.e.m. |
|---|---|---|---|---|---|---|
| $t_1$ | 0.5148 | 0.0245 | 0.0469 | 0.0038 | 0.0044 | 0.0024 |
| $t_2$ | 0.3045 | 0.0265 | 0.0515 | 0.0059 | 0.0498 | 0.0047 |
| $t_3$ | 0.3588 | 0.0268 | 0.0436 | 0.0037 | 0.0664 | 0.0177 |
| $t_4$ | 0.3128 | 0.0095 | 0.0030 | 0.0011 | 0.0043 | 0.0013 |
| $t_4$* | - | - | - | - | 0.0289 | 0.0058 |

**Supplementary Figure 5**

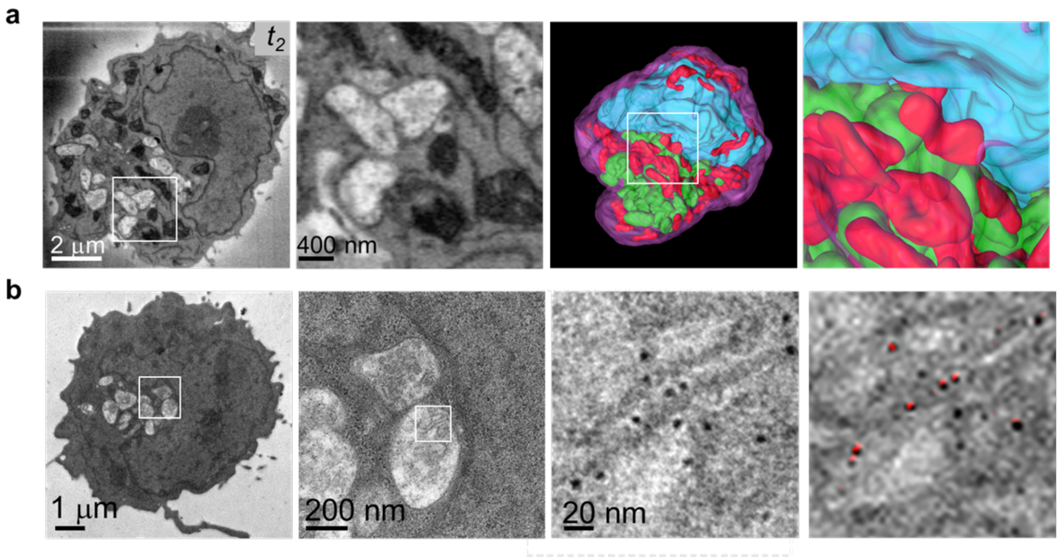


**Suppl. Figure 5.** Structural analysis of a $t_2$-stage cell. (a) SB-SEM orthoslice, showing an example of the cell analyzed, a selected magnified region containing membrane bound lysosomes, a whole cell rendered in 3D and a magnified area next to the nucleus, where the lysosomes are surrounded by mitochondria at the pole of the cell. Although 3D structural information is obtainable with SB-SEM, which is crucial in determining volumes and a general understanding of what happens on global scale of the cell, ferritin particles cannot be observed at this resolution or proven to contain iron. (b)TEM image of a thin section of a similar type of cell with an area of interest (white square), a magnified region that includes the lysosomal region (white square), a further magnified region depicting dense particulates, some of which are ferritin particles, labeled in red (confirmed with EELS), in the overlay image of the same region.

**Supplementary Figure 6**

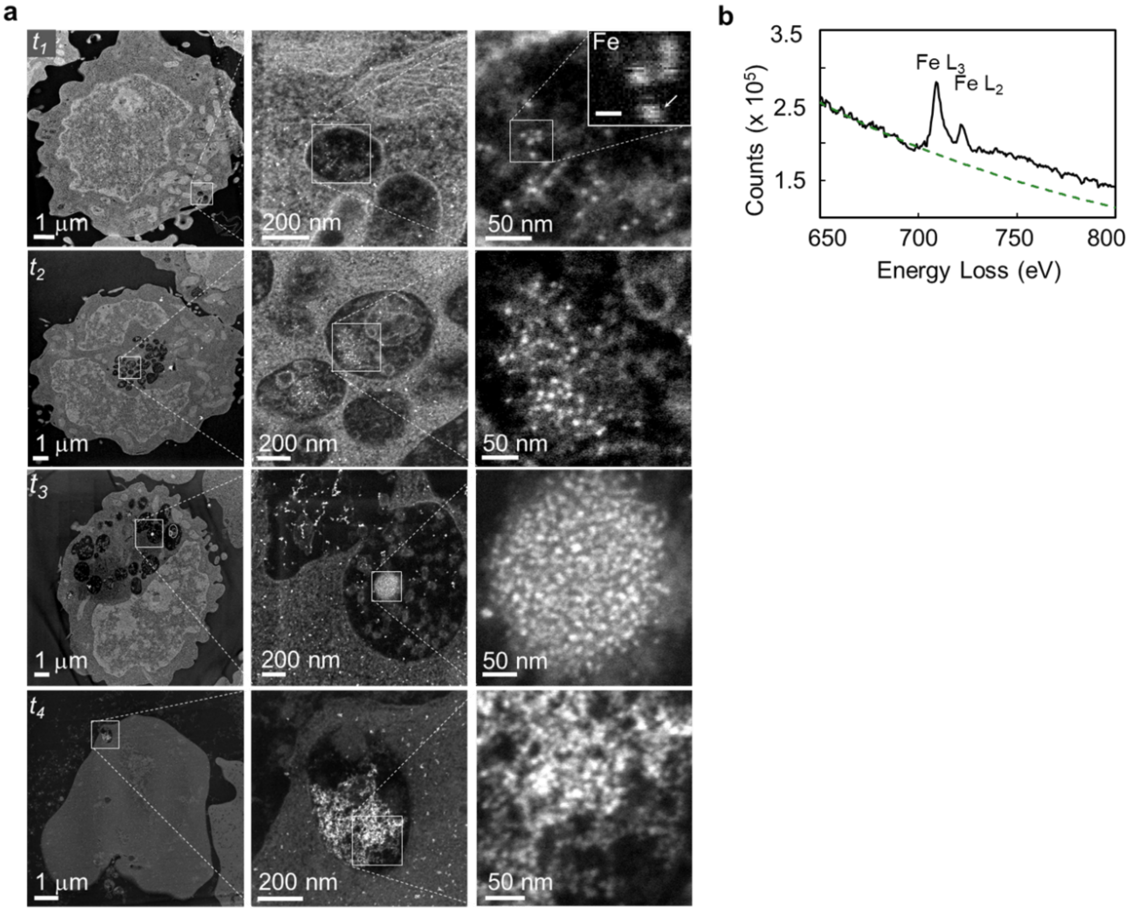


**Suppl. Figure 6.** (a) For each row of images, corresponding to $t_1$-$t_4$, a low magnification dark field STEM image (right) shows an example of a cross section of a cell with an outlined region (white square) depicting a lysosomal area, a magnified region (middle) with an outlined area (white square) and a region containing multiple ferritin particles further magnified (right). The set of images, corresponding to the $t_1$ cells, also shows an inset of the Fe map, in which individual ferritin particles are visible (micron bar = 10 nm). The intensity in this map corresponds to different amount of $Fe^{3+}$ found in those ferritin particles. (b) EELS spectra from one ferritin particle (arrow in the inset), showing the peak visible at about 710 eV corresponds to Fe $L_3$ and Fe $L_2$ edges, which is a signature of $Fe^{3+}$.

**Supplementary Note 4**

To generate distributions and their respective errors of the number of Fe atoms/cell at stages $t_1$-$t_4$ as shown in Fig. 3c, the following calculations were performed on the ADF STEM thin section images:

for $t_1$ and $t_2$ specimens, since the ferritin particles were found in most lysosomes, we used a statistical approach to obtain the total number of ferritins found in a cell $fer^{cell}$,

$$fer^{cell} = \frac{\sum fer^{lys}}{\sum A^{lys} * t} \times V^{lys} \qquad \text{(S9)}$$

in which $\sum fer^{lys}$ is the total number of ferritins found in the lysosomal region per cell in a thin section, $\sum A^{lys}$ is the total area of the lysosomes in $nm^2$ in a thin section per cell, $t$ is the section thickness in nm, which is 100 nm, and $V^{lys}$ is the total lysosomal volume in $nm^3$, a measurement that is only obtainable from segmented volume, of SB-SEM data. There were 150-200 lysosomal areas analyzed for $t_1$-$t_4$ specimens. The values used to plot the histogram in Fig. 3 are $\overline{fer}^{cell} \pm$ s.e.m.

For $t_3$ and $t_4$ stages, since most of the lysosomes did not contain ferritins and the ones that did formed stochastic clusters, we approached the analysis differently in order to propagate the error that is meaningful. First, we obtain the volume of ferritin clusters $V^{clus}$:

$$V^{clus} = \frac{\sum A^{clus}}{\sum A^{lys} * t} \times V^{lys} \qquad \text{(S10)}$$

where $\sum A^{clus}$ is the summation of all the areas that the clusters occupy in $nm^2$ in a cell of the thin section, $\sum A^{lys}$ is the total area of the lysosomes in $nm^2$ in the same cell of a thin section, $t$ is the section thickness in nm, which is 100 nm, and $V^{lys}$ is the total lysosomal volume in $nm^3$, a measurement that is only obtainable from segmented volume of SB-SEM data. From the ferritin clusters that were observed, we obtained the average number of ferritins per cluster- $fer^{clus}$. Therefore, the total number of ferritins in a cell $fer^{cell}$:

$$fer^{cell} = fer^{clus} \times V^{clus}, \tag{S11}$$

We have propagated the error for each of these operations. The area fraction's $\frac{\sum A^{clus}}{\sum A^{lys}}$ s.e.m. error was used, since there is a large lysosome to lysosome variability. As a result of these calculations the graphs Suppl. Fig. 6 are generated.

**Supplementary Figure 7**

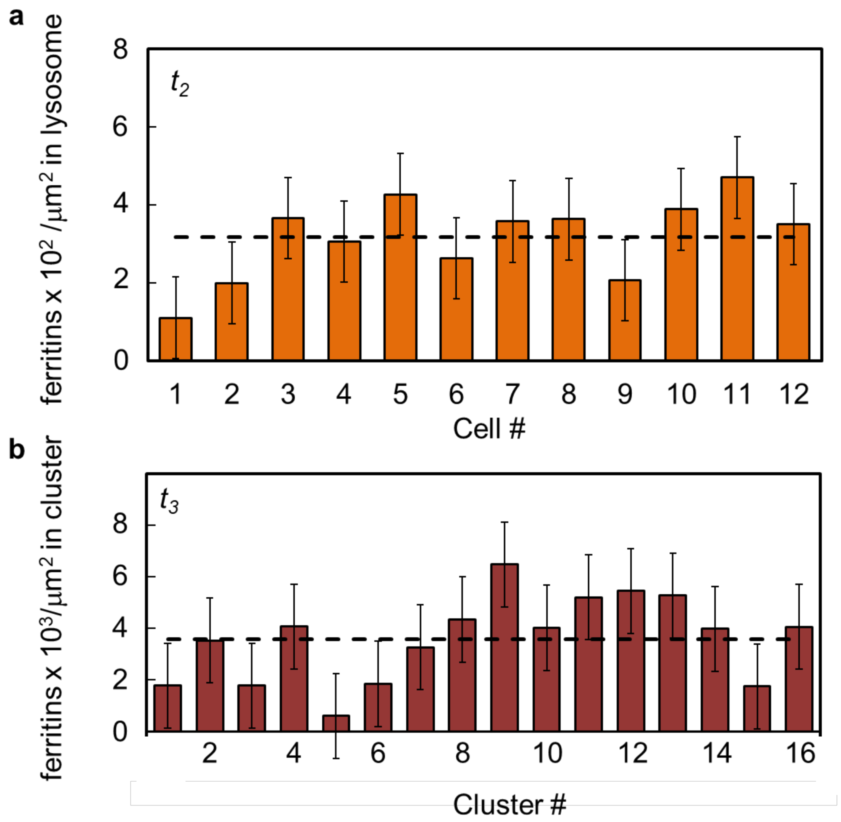


**Supplementary Fig. 7.** Counting statistics of (a) numbers of ferritins/µm$^2$ per cell in $t_2$ stage, and (b) numbers of ferritins per cluster in $t_3$ stage.

### Supplementary Figure 8

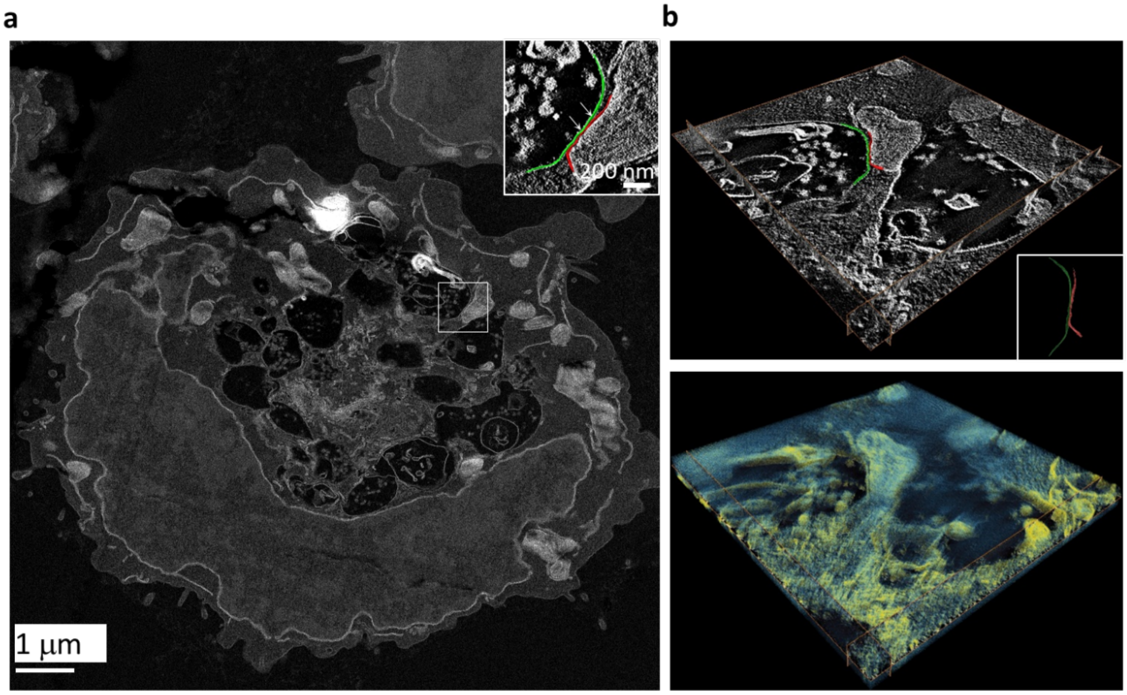


**Supplementary Fig. 8.** Dark field STEM tomography of mitochondria-lysosome contact regions of cells at $t_3$ stage. (a) Overall low magnification image of the cell showing the contact region of mitochondria and two lysosomes in the outlined square, where a tomographic tilt series was collected. Inset shows the outlined area, in which, part of the lysosomal (green) and mitochondrial (red) membranes are segmented and surface rendered. The white arrows are pointing to region where separation between the two membranes is 5-10 nm. (b, top) Orthoslice in *x-y* plane obtained from tomogram, showing the same segmented region as in inset in (a). (b, bottom) Volume-rendered tomogram, with membranes (yellow) and other structures (blue), showing close juxtaposition of lysosomal and mitochondrial membranes.

**Supplementary Figure 9**

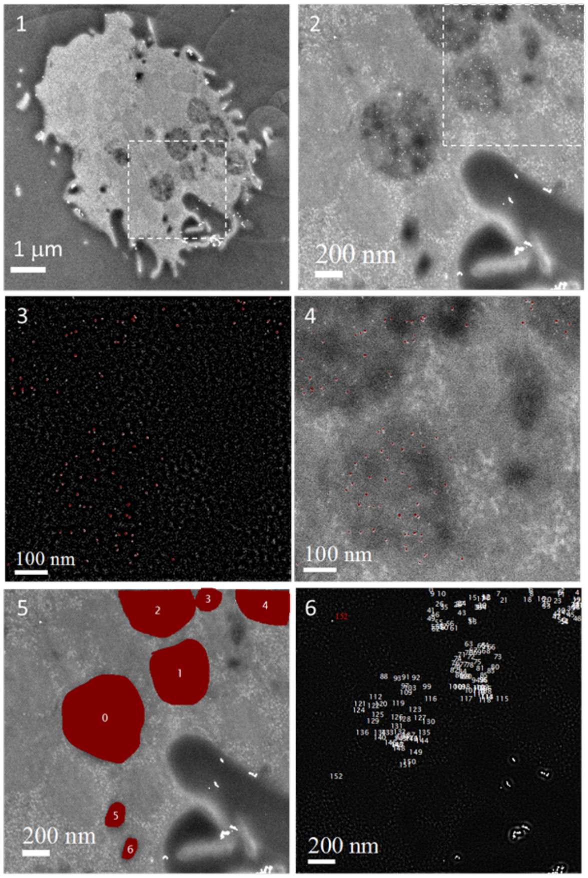


**Suppl. Figure 9.** Demonstration of the statistical approach discussed in Suppl. Note 4. (1) Dark field STEM low magnification image of a $t_2$ cell with an outlined region (white dotted square) shown in (2), which includes lysosomes with ferritins. (3) The same region as (2) with a band pass filter applied, to isolate the frequencies corresponding to the ferritin particles in the image. A mask is created and copied to the original image to check for accuracy (4). (5) A mask applied to (2) to calculate the lysosomal area $A^{lys}$. (6) The number of ferritins labeled from (2) gives $fer^{lys}$ .

**Supplementary Movie 1**

SB-SEM of a cell in the $t_2$ stage, with segmented cell membrane (magenta), nuclear membrane (cyan), mitochondria (red) and lysosomes (green).

**Supplementary Movie 2**

Dark field STEM tomogram of the mitochondria-lysosome contact region.
